# Computational and neural evidence for altered fast and slow learning from losses in gambling disorder

**DOI:** 10.1101/2024.01.08.574767

**Authors:** Kiyohito Iigaya, Tobias Larsen, Timothy Fong, John P. O’Doherty

## Abstract

Learning occurs across multiple timescales, with fast learning crucial for adapting to sudden environmental changes, and slow learning beneficial for extracting robust knowledge from multiple events. Here we asked if miscalibrated fast vs slow learning can lead to maladaptive decision-making in individuals with gambling disorder. Participants with problem gambling and a recreational gambling control group without any symptoms associated with problem gambling performed a probabilistic decision-making task involving reward-learning and loss-avoidance while being scanned with fMRI. Problem gamblers showed impaired reversal learning from losses compared to the control group, with excessive dependence on slow timescales and reduced reliance on fast timescales. fMRI data implicated the putamen, an area associated with habit, and medial prefrontal cortex (PFC) in slow loss-value encoding, with significantly more robust encoding in medial PFC in problem gamblers compared to controls. Problem gamblers also exhibited stronger loss prediction error encoding in the insular cortex. These findings suggest that individuals with problem gambling have an impaired ability to adjust their predictions following losses, manifested by a stronger influence of slow value learning. This impairment could contribute to the behavioral inflexibility of problem gamblers, particularly the persistence in gambling behavior typically observed in those individuals after incurring loss outcomes.

## Introduction

The world changes across various timescales (e.g., the temperature changes both daily and seasonally). Hence, humans and animals must have developed sophisticated mechanisms to learn and adapt over such timescales. Extensive research has shown behavioral and neural evidence for learning across different timescales.^1–20^ While such multi-timescale learning is generally effective, improper weighing of information from different timescales can lead to maladaptive behaviors. For instance, over-reliance on slow learning in rapidly changing environments may result in maladaptive decision-making.^13, 21^ This raises an intriguing question: could such miscalibrated learning be relevant to psychiatric conditions, such as gambling disorder, and if so, what neural mechanisms underlie this phenomenon?

Gambling disorder (GD) is characterized by persistent and problematic gambling behavior that often leads to significant impairment and distress.^22, 23^ A key behavioral symptom of gambling disorder is loss chasing, whereby individuals persist in gambling behavior ostensibly to remediate their financial position after extensive losses have already been incurred.^24, 25^

The underlying neurocomputational basis of the propensity to chase losses in gambling disorder is currently unclear. A number of studies have utilized behavioral economics methods to probe attitudes to loss in gambling disorder during choice behavior, in which one natural hypothesis could be that individuals with gambling disorder have reduced loss aversion. However, collectively such studies have found varying and somewhat inconsistent evidence for behavioral differences between gamblers and non-gamblers in the degree of loss aversion during choice.^26–29^ Another direction of research has been to test for the sensitivity of individuals with gambling disorder to changing their behavior following feedback. One notable behavioral finding that aligns with persistent behavior in gamblers, especially in the face of losses, is increased perseveration on reversal learning paradigms.^30, 31^ Several recent studies have utilized reinforcement-learning models to examine altered computational substrates for learning in gambling disorder, and these studies have identified a number of differences in reinforcement-learning computations, such as decreased direct exploration,^32^ as well as altered learning rates for both gains and losses^33^

The aim of the present study was to examine the role of alterations in slow and fast learning in gambling disorder. We utilized a variant of a task designed to probe reinforcement-learning in both gain and loss contexts separately.^34^ Specifically, in the gain condition, participants could choose between two stimuli associated with differing probabilities of winning a monetary reward outcome. Their goal in that condition was to learn to choose the stimulus associated with the highest probability of winning money. In the loss avoidance condition, participants had to choose between two stimuli associated with differing probabilities of obtaining a monetary loss. Their goal in that condition was to choose the stimulus associated with the lowest probability of losing. After a variable number of trials, the contingencies were reversed in both conditions so that the stimuli previously associated with the highest probability of winning, or the lowest probability of losing were no longer advantageous, thus necessitating the need for participants to their choice of stimuli in each condition.

Utilizing a formal computational model-based approach, we differentiated between the contribution of two different forms of reinforcement-learning: slow and fast learning in guiding behavioral performance on this task. We could further separately evaluate the contribution of these two processes in learning about gains and in learning about losses. We hypothesized that individuals with gambling disorder might rely excessively on slow-learning when making decisions, as an overemphasis on slow-learning processes could provide an explanation for the behavioral phenomenon of increased perseveration previously reported in such individuals. Furthermore, we aimed to test whether a reliance on increased slow-learning would be especially prominent when learning from losses, which could potentially account more specifically for the behavioral observation of increased loss chasing in gamblers. Increased reliance on slow-learning from losses could lead to the behavioral manifestation of a reduced tendency to switch away from a losing option, because changes in the expectation about the future value of that option would be updated only very slowly, even after accumulating losses.

To examine this hypothesis, we scanned 20 individuals who met the diagnostic criteria for gambling disorder using fMRI while they performed the gain/loss learning task, and we also scanned 20 recreational gamblers as a well-matched comparison group (who did not reach the diagnostic criteria of gambling disorder). Participants were recruited via flyers from the Los Angeles area. The study was conducted in a double-blinded manner, in which both the experimenter conducting the fMRI study and the participants themselves were unaware of the diagnostic category of the participants during the data acquisition.

In addition to testing our computational hypothesis at the behavioral level, we also aimed to examine the extent to which neural responses related to slow learning were distinct in individuals with gambling disorder, in order to help pinpoint the specific neural computational substrates associated with altered reinforcement learning in this cohort. We hypothesized that brain regions previously implicated in loss-related learning and aversion-related processing, such as the insular cortex and striatum^35–38^ would show alterations in the neural correlates of loss-related computations in individuals with gambling disorder.

## Results

### The study design

After an initial phone screening, we conducted a clinical interview of 110 participants who reported engaging in regular gambling but who had no prior psychiatric diagnoses, including gambling disorder (GD). From this initial cohort, we identified 20 individuals who met the diagnostic criteria for gambling disorder (referred to as the GD group) and 20 “recreational” gamblers (who had at least some recent experience of having gambled within the past 6 months, but who had none of the diagnostic criteria of gambling disorder; referred to as the healthy volunteer [HV] group). Importantly, this study utilized a “double-blind” procedure, ensuring that neither the experimenter running the fMRI study nor the participants being scanned were aware of the outcome of the clinical assessment during the administration of the study protocol.

During the study, participants underwent fMRI scanning while performing an adaptation of a well-established probabilistic reward and avoidance learning task^34^ (**Figure 1A**). On each trial, participants were presented with two alternative target stimuli on the screen and were required to make a choice between them. The stimuli were organized into three sets, with each pair belonging to one of three conditions. In the reward condition, the stimuli were associated with the chance of receiving monetary rewards. In the loss avoidance condition, the stimuli were associated with the chance of avoiding monetary losses. The neutral condition consisted of stimuli with no associated outcomes.

**Figure 1.**
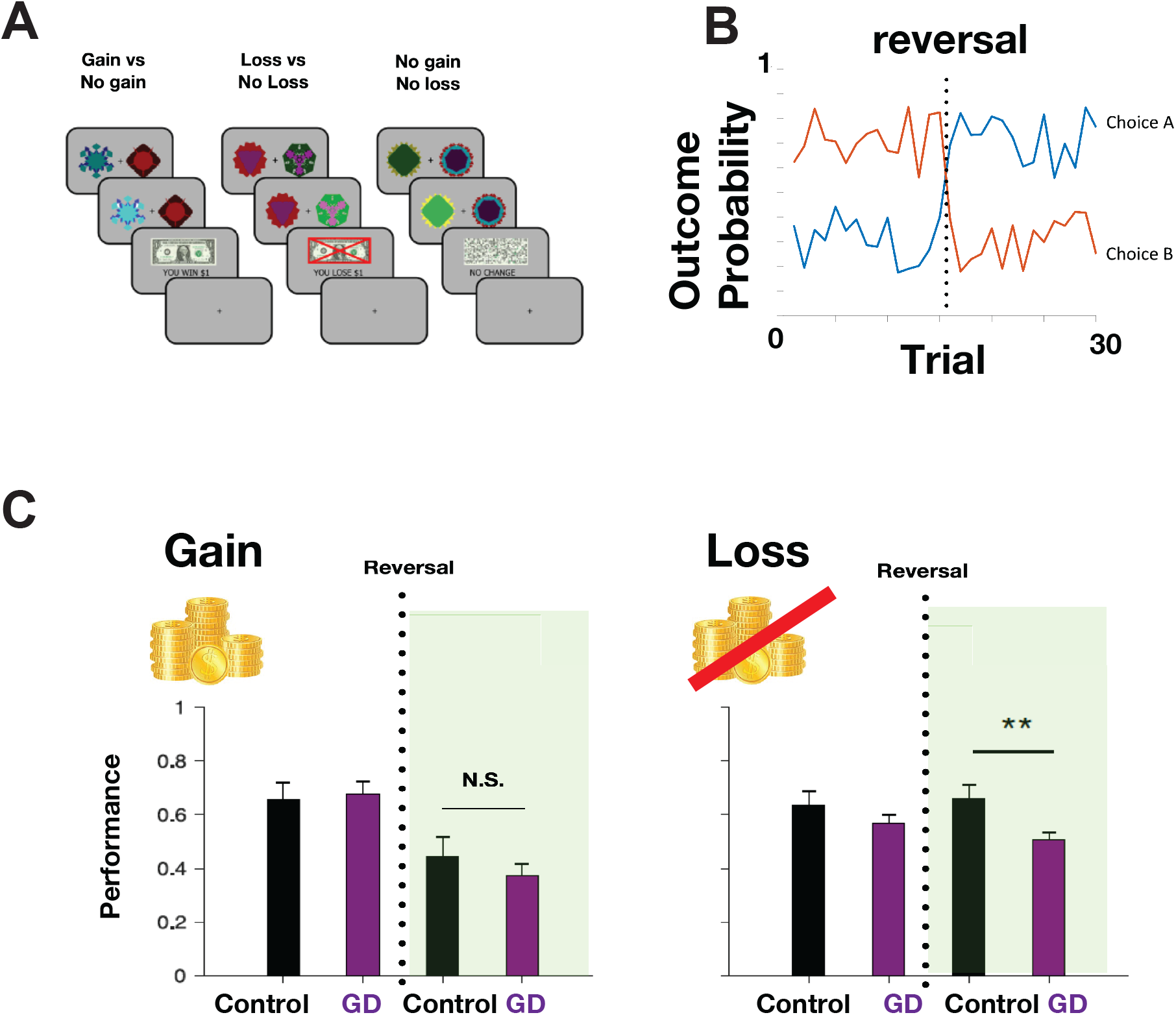
Task and behavior (**A**). The task. On each trial, a participant was presented with two stimuli, each of which was associated with a unique probability of outcomes. There were three trial types. On trials in the gain condition, the stimuli were associated with potential monetary gain outcomes. On trials in the loss-avoidance condition, the stimuli were associated with potential monetary losses. On trials in the neutral trials, participants received no monetary outcomes. (**B**). Example outcome probabilities in the reward condition. The probabilities of reward associated with the two stimuli (choice A and choice B) are plotted as a function of the trial number. Choice B is initially more rewarding than choice A. After about 15 trials, the reward probabilities are reversed. (**C**). Behavioral performance, measured by the number of choices allocated to the target with a higher reward (or a lower loss) probability divided by the number of trials, is shown before and after the reversal points for control and individuals with gambling disorder. In the gain condition, there was no significant difference between groups before or after the reversal. However, the difference between before and after reversal within each group was significant (*p <* 0.01 permutation test for each group). In the loss avoidance condition, there was a significant difference in performance between groups after the reversal (*p <* 0.01 permutation test), but no significant difference between the groups before the reversal.

The outcome probabilities remained relatively stable for approximately 15 trials in each condition. Subsequently, without any warning, the outcome probabilities were reversed, challenging participants to adapt their choices based on their previous outcome experiences (**Figure 1B**). Each condition underwent one reversal, and a total of 30 trials per condition were conducted across two sessions.

### Individuals with gambling disorder showed impaired reversal learning in the loss-avoidance condition

We first investigated if there was any difference in behavioral performance on the task between PG and HV. As seen in **Figure 1C**, we found that there was no significant difference between groups before or after the contingency change in the gain condition. However, we found that the GD group performed significantly worse than the HVs in the loss avoidance condition on trials after the reversal (*p <* 0.01 permutation test). This suggests that individuals with GD had impaired reversal learning from loss outcomes compared to the HV group. Both groups performed significantly worse after the reversal than before the reversal in the gain condition (*p <* 0.01 permutation test for each group), though in that condition, there was no significant difference between the groups (permutation test).

### Computational modeling analysis suggests over-weighting of slow learning relative to fast learning in gambling disorder

We then performed a computational model-driven analysis to examine if the observed behavioral performance differences could be captured by over-weighting of slowly learned value. We used a variant of a previously validated multi-timescale learning model of decision making.^12, 13, 39, 40^ The model integrates reward history across two timescales (fast and slow) and computes the value of choice (**Figure 2A**). The model has free parameters that determine the impact of fast and slow learning (*w*_*fast*_ / *w*_*slow*_; please see the Methods section for the full model description).

**Figure 2.**
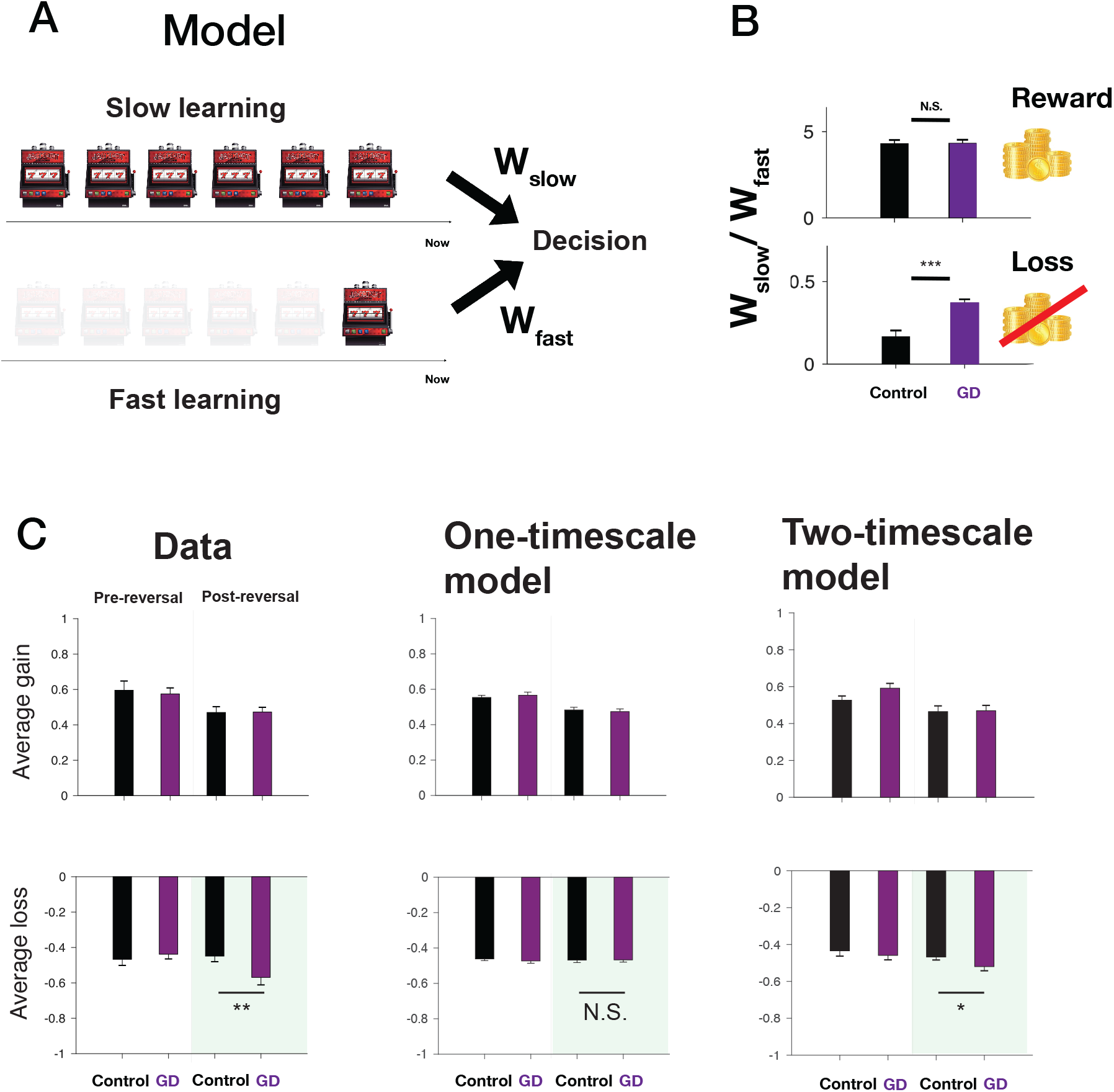
A computational model describing learning of reward history across multiple timescales captures human behavior on the task. (**A**). Schematic of the computational model. Gain or loss history is integrated over two timescales independently to compute fast and slow values. The two values are then weighted with relative weights to compute an overall decision-value for each condition separately. (**B**). The model’s estimates of relative weights. The relative weights assigned to slowly learned value with respect to the fast value are plotted for the HV and PG groups for reward trials (top) and loss trials (loss). There is no significant difference between groups in gain trials, but the individuals with gambling disorder show significantly larger weight than the control in loss trials (*p <* 0.001 permutation test). (**C**). Model simulations show that the classic reinforcement learning model does not capture data, but the two-timescale model does. From left to right: data, one-timescale model simulation, and two-timescale model simulation. The average monetary gain received per trial is shown before and after reversals for the HV (black) and GD (purple). The data show significant differences in behavior between groups after reversal in loss trials. This is not captured by the standard RL model (middle), but the two-timescale learning model captured this (right).

We performed a model fitting using a hierarchical Bayesian method.^12^ We found that the relative impact of fast and slow learning (*w*_*fast*_ / *w*_*slow*_) was similar for the PG and HV groups in the reward condition. However, the impact of slow learning compared to fast learning was significantly greater for the PG group in the loss-avoidance condition compared to the HV group (**Figure 2B**). This confirms our hypothesis.

To establish the validity of our two-time scale model, we also fit a standard reinforcement learning model with a single learning rate to the behavioral task data across groups. When comparing the fit of this standard model to the data against that of the two-time scale model, the integrated-BIC score^12, 41^ for the standard RL was 4533, and the one for the two-timescale model was 4383 in the logarithmic scale, implying that the two-timescale model significantly outperformed the standard RL with a single timescale. Further, model simulations showed that the standard single learning rate model fails to capture the behavioral signature in the real human choice data, namely the group differences we found in performance after reversals on loss trials. The two-timescale model, on the other hand, captures this core behavioral feature well (**Figure 2C**).

### fMRI reveals neural correlates of slowly learned value in the putamen and the ACC/dmPFC

Behavioral analyses suggest that the key difference between control and GD is the slow value computed on loss trials. The next question, therefore, is to determine how slow value is computed in the brain, with a particular focus on the loss avoidance condition, before investigating neural differences between groups in such value signals. For this, we performed a standard GLM analysis utilizing parametric regressors with variables from the two-time scale computational model (see methods). For each participant, we entered parametric regressors generated using individual subject-level parameter estimates (maximum a posteriori probability, MAP, estimates) drawn from the hierarchical behavioral model fitting procedure (see methods).

We first asked how the slow value is represented at the time of every trial onset in the loss avoidance condition, using all participants’ data (i.e., pooling over the two groups). A whole brain analysis with cluster-level FWE error correction revealed that slow value from the computational model was correlated with activity in the left putamen (peak coordinate [-24,-2,0]) and anterior cingulate cortex (peak coordinate [-6,44,28]) (**Figure 3A**).

**Figure 3.**
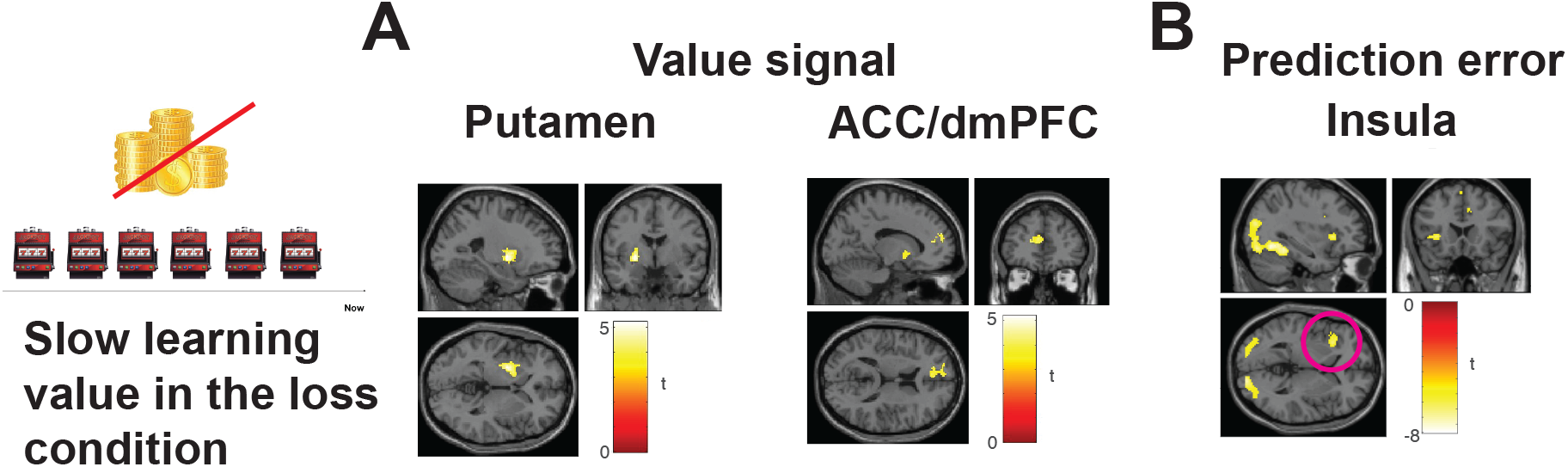
fMRI correlates of the slowly learned value component from the two time-scale computational model. (**A**). A cluster of voxels in the left putamen is significantly correlated with the model’s predicted signal (whole-brain cFWE *p <* 0.05 with height threshold at *p <* 0.001. This result comes from an analysis that pooled across the two groups. In addition, a cluster of voxels in the ACC is significantly correlated with the model’s predicted signal (whole-brain cFWE *p <* 0.05 with height threshold at *p <* 0.001. This result comes from an analysis that pooled across the two groups. (**B**). A cluster in the left insula was significantly correlated with prediction error signals from the slow-learning component of the two-time-step model (whole-brain cFWE *p <* 0.05 with height threshold at *p <* 0.001.

In order to gain insight into how slow-value signals are computed on the basis of trial and error feedback (loss or no-loss), we next examined how prediction error signals are represented at the timing of outcome presentation. Aversive prediction error signals were generated using the model with the MAP estimates of parameters for each trial for each participant. A whole brain analysis with cluster-level FWE correction revealed that the left insula is significantly negatively correlated with an aversive-learning prediction error signal (**Figure 3B**). This insular PE signal responds positively with unexpected losses and negatively with unexpected loss omission, across all participants.

We also examined fast value signals generated from the model in the loss condition within the same GLM analysis. We found no significant activity correlating with fast-value at whole brain FWE corrected levels. However, at an uncorrected threshold, a cluster of voxels in mPFC was found to correlate the fast value signal (see **Table 1**).

**Table 1:** Non-significant fMRI results. Peak voxels and the t-values for different contrasts are shown.

| Contrast | Region | (x,y,z) | T |
| --- | --- | --- | --- |
| Fast Negative | dmPFC | (-14,50,28) | -3.52 |
| Slow Positive | dmPFC | (6,60,32) | 3.49 |
| Fast Positive | vmPFC | (6,30,-6) | -3.3 |

Finally, we also tested for BOLD correlates of slow and fast values in the gain condition in the same GLM analysis. We found no clusters surviving whole-brain FWE correction for these contrasts. However, a cluster in the dmPFC was found to correlate with slow value in gain trials at an uncorrected threshold (**Table 1**). A cluster in vmPFC was found to be correlated with fast value in gain trials at an uncorrected threshold (**Table 1**)

### fMRI reveals different neural responses in GD group compared to HV group in dACC and insular cortex

We next tested for differences between the GD and HV groups in the fMRI activity related to slow and fast-value in the loss condition, a particular focus for our fMRI analysis given the behavioral findings that the GD group is different from HV when learning from losses but not gains. For this, we utilized regions of interest defined on the basis of the clusters identified from the pooled group results. Note that selecting ROIs based on the pooled group results to investigate group differences is justified on the basis that such an ROI analysis identifies relevant voxels responsive to the task variables, while such an ROI selection strategy does not positively bias the likelihood of observing a significant group difference (see **Methods** for a description of simulations that demonstrate this).

Specifically, to test whether correlations with slow learning are greater for the GD compared to the HV groups, we used regions of interest (ROI) defined from the putamen and anterior cingulate clusters identified from the pooled analysis. We found no significant difference in the strength of the correlation with slow-value between groups in the putamen ROI, but the correlation with slow-value in the ACC ROI was significantly greater in the GD than HV group (**Figure 4A**).

**Figure 4.**
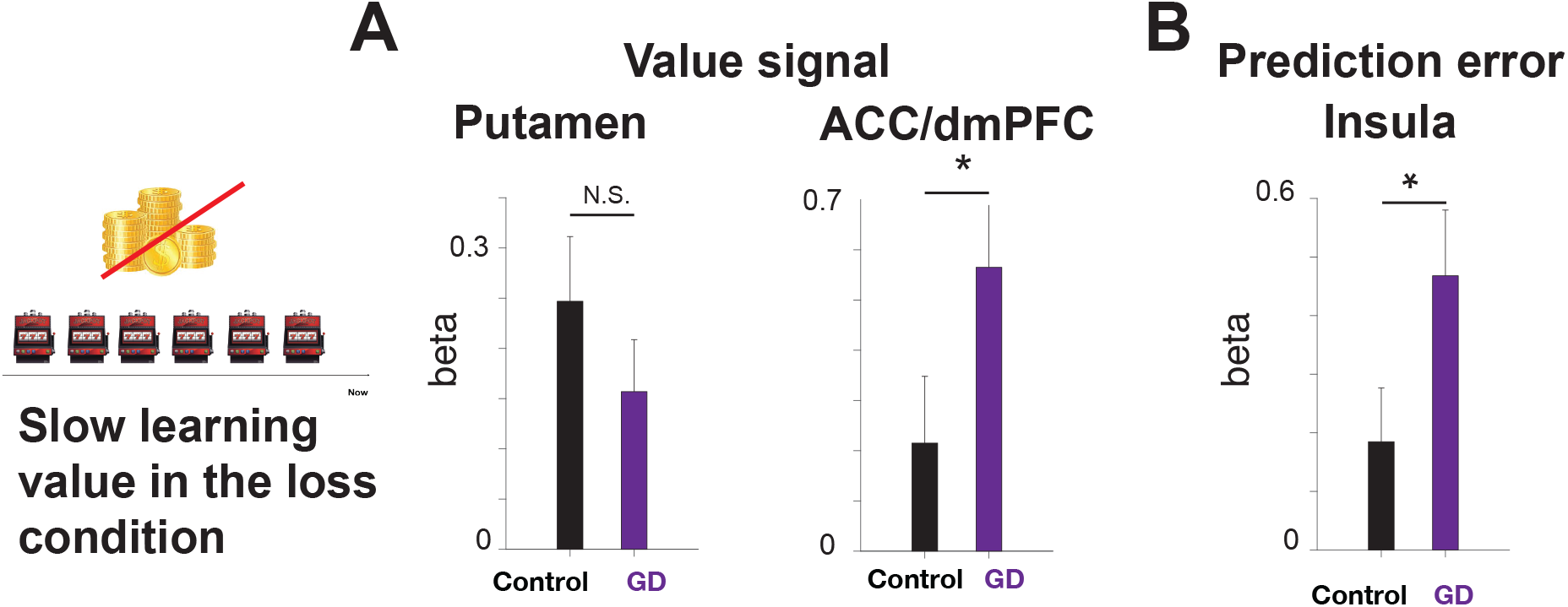
Group difference in the fMRI correlates of slow-value learning in loss trials. (**A**). The slow value signal. There was no significant difference between groups in an ROI defined on the putamen cluster identified from the pooled analysis. However, a cluster of voxels in the ACC is significantly correlated with the model’s predicted signal (whole-brain cFWE *p <* 0.05 with height threshold at *p <* 0.001. This result comes from an analysis that pooled across the two groups. (**B**). The prediction error signal. The magnitude of correlation was found to be significantly greater for the GD than HV groups (*p <* 0.05 permutation test).

Next, we tested for a difference in prediction error responses related to slow learning. Specifically, we asked whether GDs or HVs showed a stronger or weaker correlation with aversive-learning prediction errors in the left insular cortex ROI defined from the cluster identified via the pooled analysis. We found that individuals with GD showed a significantly greater correlation with aversive prediction errors in loss trials compared to the HV group (**Figure 4B**).

For completeness, we also tested for differences in the neural responses between the groups related to fast value during the loss condition, focusing on the cluster that was identified in mPFC from the pooled analysis. No significant difference was found between groups in this cluster.

We also tested for group differences related to slow-value in the gain condition, focusing on the dmPFC cluster reported in the pooled analysis. We found no significant difference between groups in that signal.

Lastly, we tested for a difference between groups in the vmPFC cluster identified as responding to fast-value during the gain condition in the pooled analysis. The correlation with fast-value in that cluster was significantly stronger for the control group than the GD group (*p <* 0.05). However, the correlation with fast value was negative, meaning that trials after no reward show stronger activation than trials after reward (**Table 1**).

## Discussion

We examined behavioral and neural differences in reinforcement-learning mechanisms pertaining to gain-seeking and loss avoidance between recreational gamblers who did not reach any of the diagnostic criteria for gambling disorder (HV group) and individuals with gambling disorder (GD group). Our hypothesis was that individuals with GD would exhibit stronger learning impairments from recent losses due to an increased reliance on slow learning. We found evidence supporting this hypothesis through behavioral analyses, where individuals with GD showed impaired performance after the reversal of the loss-probability association between choices, indicative of their reliance on slowly learned values. We further confirmed our hypothesis using computational modeling of multiple timescale learning.

Impaired behavioral adjustments following loss in gambling disorder have been well-documented. A notable phenomenon is loss chasing,^25, 42, 43^ where gamblers amplify their betting after losses. Our results are consistent with loss-chasing, and our model captures this as a result of ignoring recent losses due to an over-reliance on slow learning.

To delve into the neural mechanisms underlying these behavioral differences, we conducted a model-based fMRI analysis. Our results revealed correlations of slow value during the loss avoidance condition in both the putamen and ACC. Previous studies in healthy volunteers have implicated the putamen in habits.^44, 45^ The present findings implicating the putamen as tracking slowly-learned value signals could align with those prior results. It has been suggested that model-free reinforcement-learning provides a computational account for habit learning.^46, 47^ However, the precise relationship between model-free learning and habits has remained unclear at both behavioral and neural levels.^48–50^

Here, we have dissociated two distinct components of reinforcement learning: slow and fast learning. Both of these forms of learning, as described here, can be considered to be model-free. However, the present neural findings relating slow-value to the posterior putamen raise the possibility that the relationship between habits and model-free learning might be more closely aligned with slow as opposed to fast model-free learning components. One complication for this interpretation of our findings is that we observed slow-value signals in the putamen in the loss avoidance condition and not in the gain condition. One possible account for these findings, is that habitization might play a stronger role early on in loss avoidance than in gain-learning, thereby more rapidly engaging the putamen. Further research will be necessary to compare the emergence of habitization under appetitive and aversive learning conditions. However, in any event, the putamen did not differ significantly between the GD and HV groups, suggesting that this brain structure does not directly account for the impaired loss-learning observed in individuals with gambling disorder.

In addition to the region of posterior putamen aligned with slow-value learning, we also found slow-value learning signals in the anterior cingulate cortex. Crucially, the anterior cingulate cortex slow-value areas did show significant differences between the gambling disorder and recreational gambling group, in the direction of enhanced slow-learning signals in the gambling disorder group. The ACC has been suggested to play a central role in integrating various information for the purpose of computing decision values,^51^ which in the present study could reflect the fact that the GD group exhibits an increased reliance on utilizing slowly-learned values to guide their choices.

One possible computational mechanism by which enhanced slow-value components might emerge in the first place in the gambling disorder group is via alterations in the means by which value signals are learned. In typical reinforcement-learning accounts, this is assumed to occur via prediction errors that signal the difference in value between expected and actual outcomes. Here, we examined prediction errors that are involved in slow-learning. During the loss avoidance condition, we tested for such prediction error signals in the brain, and we found a slow-learning aversive prediction error to be encoded in the left insular cortex. The insular cortex has previously been suggested to encode aversive learning prediction errors on a similar task.^34^ This insular cortex signal was found to be significantly enhanced in the gambling disorder group compared to the recreational gambling group. This suggests that individuals with gambling disorder may be experiencing enhanced slow-learning contributions to behavior, because of an enhanced engagement of slow-learning prediction error signals during the reinforcement-learning process. The identification of alterations in prediction error coding in the anterior insula in the gambling disorder group, aligns with previous studies that have implicated the insula in gambling disorder.^33, 52–56^ The insula has also been implicated in other addiction disorders, such as substance use disorders in general,^57^ as well as in the encoding of risk more generally.^58^ Our findings suggest a specific computational role, especially in GD, in the form of prediction error from losses, related to the slow-learning component.

In light of our behavioral findings that the individuals with gambling disorder were impaired when learning from losses, we focused the neuroimaging analysis on the loss avoidance condition. However, we did also find some tentative evidence for altered responses in gambling disorder during gain-learning, specifically in the ventromedial prefrontal cortex. Thus, it may be premature to conclude that individuals with gambling disorder are exclusively impaired in the loss processing domain. Indeed, numerous fMRI studies of gambling disorder have reported alterations in reward-related responses in the medial prefrontal cortex and ventral striatum^56, 59–62^ However, it seems reasonable to conclude that alterations in slow-learning have the potential to account for an important element of real-world behavior in individuals with gambling disorder, whereby they persist in gambling even after sustaining substantial monetary losses.

A limitation of the present study is that we utilized a relatively small sample size of 20 individuals in each group. To assess the robustness of the present findings, it will therefore be important to follow up with a replication study in a larger cohort. Despite this limitation, it is important to note that the present study has a number of significant methodological advantages over the typical approach utilized in the gambling disorder literature. First of all, unlike studies that recruit patients with gambling disorder from a gambling disorder clinic, here we recruited participants with gambling disorder directly from the community. While the clinic provides a convenient means of recruitment, it has several downsides. These include the fact that the subset of individuals with gambling disorder actively seeking treatment may be qualitatively distinct in a number of respects from those who are not actively seeking treatment, thereby introducing a potential sampling bias. A further concern is that if such patients have already been in treatment through the clinic (as would be the case for many such patients), then any assessment of their behavior and/or neural responses will be confounded by treatment effects. The individuals in our study indicated that they were not actively seeking treatment (though we did provide them with information about how to do so after they participated in our study). Secondly, we recruited our comparison group in the same way as the individuals with gambling disorder, and we ensured they had all recently gambled recreationally while showing no clinical signs of the disorder. This ensured that our healthy comparison group was well-matched demographically. Finally, to further minimize bias, we implemented a rigorous double-blind experimental design. Neither the participants nor the experimenters were aware of the participants’ classification as GD or control. Although more time-consuming than traditional designs, this approach allowed us to eliminate experimenter demand biases arising from knowing the participants’ diagnoses. Consequently, in spite of our limited sample size, the present study improves in a number of other important methodological respects on the typical approach used in the field on account of the careful recruitment and design procedures we employed.

More generally, the present study provides an example of how careful computational modeling can, when applied to behavioral and fMRI data, provide insight into the nature of the computations that may be altered in psychiatric disease.^63–66^ The findings of enhanced neural responses in the anterior cingulate and anterior insular related to slow-value encoding and learning, suggest the possibility that these neural responses could be precisely targeted for treatment in individuals with gambling disorder, such as by, for example, through focused transcranial magnetic stimulation.^67^

## Methods

### Participants

We recruited 20 participants into our gambling disorder group (GD) and 20 participants into a non-pathological gambling healthy volunteer group (HV). Participants were recruited from the greater Los Angeles area using a combination of posted flyers and Craigslist. A requirement for participation in the study was that participants had gambled at least once in the last six months. We recruited people from the population at large as opposed to targeting individuals already in treatment for pathological gambling as we aimed to avoid the potential confounding effects of individuals who have already decided to seek treatment potentially being different from the population of pathological gamblers at large, as well as avoiding any potential confounding effects of the treatment program itself on our measured effects. We further aimed to specifically recruit “recreational” gamblers into our HV group, that is, people who have at least some experience of gambling (and have recently gambled in the past six months), but who do not reach the clinical threshold for problem gambling. We reasoned that these individuals would be better matched to our PG group in terms of life experience, demographics, and exposure to gambling-related stimuli and environments than individuals with absolutely no gambling experience whatsoever.

More than 400 potential participants replied to our advertisement and were subjected to an initial telephone screening to determine their eligibility. 110 passed the initial screening and were subsequently invited to come to the lab to participate in a more detailed in-person clinical interview. The participants assigned to the problem gambling group were selected to have a score of 2 or more on the clinical scale in DSM-IV, while those assigned to the control group all had a score of 0. Note that data collection for this study was begun in 2013, prior to the official release of DSM-5; hence, we utilized the diagnostic criteria for DSM-IV throughout. However, given the participant’s characteristics, we don’t expect that group assignment would have changed had the study been implemented under DSM-5 criteria. The participants in both groups were furthermore required to be free of axis I disorders (e.g., anxiety, depression, PTSD, as well as schizophrenia) and were not currently taking any neuromodulatory medication.

We continued recruitment until we reached our target goal of 20 participants in the problem gambling group and 20 matched controls in the recreational gambling group. None of the participants in our final sample in either group reported being previously diagnosed with problem gambling, and none reported seeking treatment for gambling-related problems. The recruitment and experimental administration were organized in a double-blind fashion so that the researcher responsible for running the fMRI experiment was not aware of the participant’s group status until after the data were collected. One additional participant (who would have been assigned to the PG group) was interviewed and scanned but was subsequently excluded due to an incidental finding on the structural scan.

The participants were remunerated $20-30 for participating in the interview session and $50-65 for participating in the scanning session. All participants were also given information about resources available for problem gambling, although they were not informed of the outcome of the clinical interview. The research was approved by the Institutional Review Boards of the California Institute of Technology and the University of California, Los Angeles, and each participant gave informed consent.

The participants were well matched on demographic variables, including age, years of education, and gender. The PG group consisted of 9 females and 11 males with an average age of 37.9 (sd 12.3) and an average of 13.6 (sd 4.8) years of education. The HV group consisted of 20 participants (10 female) with an average age of 36.9 (sd 11.6) and an average of 14.6 (sd 3.9) years of education.

### Task description

The task performed by the participants was a modified version of the reward and avoidance learning task first used in.^34^ The task involved two-alternative free choices with three intermixed conditions: a reward, a loss avoidance, and a neutral condition. On each trial, the participant is presented with one of three pairs of fractal stimuli. Each pair is specific to one of the three conditions (Fig. 1). On each trial, the participant then has to choose between the two presented fractals, each of which is associated with differing probabilities of yielding particular outcomes. In the reward condition, participants could either win $1, or else gain nothing, while in the loss avoidance condition, participants could either lose $1 or else gain nothing. The neutral condition probabilistically involves the visual feedback of a scrambled dollar bill or else no outcome, which in either case results in no change in overall winnings. Irrespective of the condition, when the trial yields no outcome, no feedback is given, and the fixation cross of the next trial is presented.

An experimental session consists of 2 blocks of 90 trials (30 trials per condition), with different sets of stimuli in each block. On each trial, one of the stimuli has a 70% chance of yielding an outcome, and the other has a 30% chance. After 10-20 trials (independently jittered between conditions), the outcome probabilities are reversed within each condition.

Before the experiment, participants are informed that the three conditions exist, but not which stimuli are associated with each particular condition. They are furthermore informed that the amount they earn during the experiment will be added to a base pay of $40, so that they could potentially either win more than $40, or else come away with less than $40 depending on how much money they won or lost while performing the task. Thus, in the reward condition, participants should aim to choose the fractal associated with the greatest probability of winning $1, thereby adding to their total winnings, while in the loss condition, they should aim to choose the option associated with the lowest probability of losing $1, thereby avoiding decrementing their overall winnings.

### fMRI data acquisition

The fMRI data were acquired using a Siemens Tim Trio 3T scanner located at the Caltech Brain Imaging Center. For each block of trials, 185 volumes of 44 slices covering the whole brain were recorded interleaved ascending at a 30-degree angle, TR = 2.78s, TE=30ms, voxel size 3mm isotropic.

### Computational modeling

We aimed to contrast a standard reinforcement learning model that utilized a single learning rate with a two-timescale model that learns reward history over two distinct timescales. For this purpose, we used a simple Q-learning algorithm with a single learning rate alongside a model that learns reward history across multiple timescales that has been previously validated. The spirit of the two timescale model is that it captures a combination of win-stay, lose-switch type of behavior and slower reinforcement learning.

The model assumes that the value of stimuli is computed as a sum of the values learned both quickly and slowly.

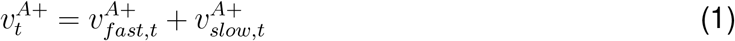

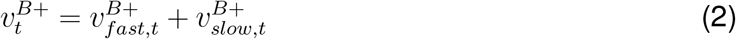

where 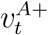 is the value of stimulus 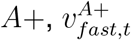 is the value of *A*+ on trial *t*, learned over a short timescale and 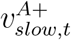 is the one learned over a longer timescale. Here, we express *A*+ and *B*+ as two choice targets presented in gain trials. In principle, the fast and slow values follow the same learning rule but with different parameters:

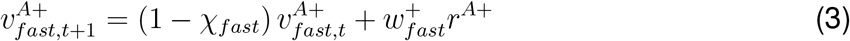

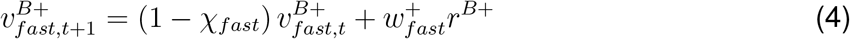

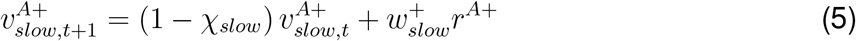

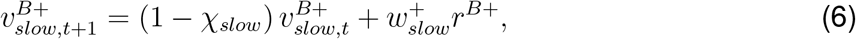

where *χ*_*fast*_ (*χ*_*slow*_) is the learning (or forgetting) rate in the fast (slow) system and 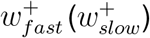 determines the impact of outcome to the value. *r*^*A*+^ = 1 (*r*^*R*+^ = 1) if a reward is obtained from *A*+ (*B*+) on trial t, or 0 otherwise. These equations can be re-written (for example, for the slow values) as

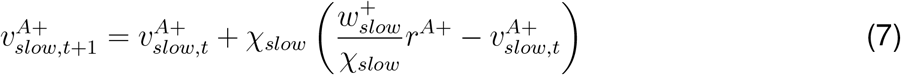

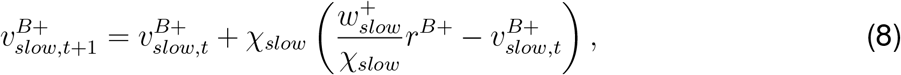

meaning that these are equivalent to a forgetful Q-learning rule^68, 69^ with a learning rate *χ*_*slow*_ and reward sensitivity 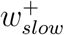. Thus the prediction error can be expressed as:

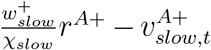 for *A*+ and 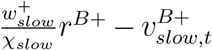 for *B*+. The choice probability is expressed as

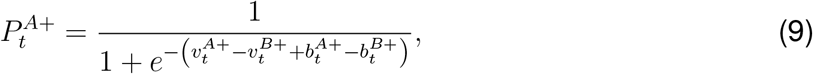

where 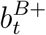 is a bias from the previous choice (choice kernel):

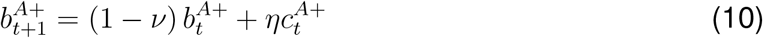

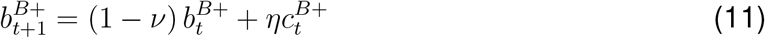

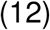

with *ν* decay constant and *η* is the weight and 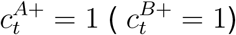 if *A*+ (*B*+) is chosen on trial t while 0 otherwise.

Because we have only 30 trials per participant, fitting this full model is not possible. Therefore we simplified the above model by assuming the time constant of the fast system to be one trial, or *χ*_*fast*_ = 1 and *ν* = 1. Note that *η*, is normally negative,^12, 70, 71^ meaning that the choice kernel normally captures a tendency towards alternation. The combination of fast reward value and fast choice kernel produces behavior similar to win-stay, lose-switch behavior.

### Model fitting

In order to determine the distribution of model parameters **h**, we conducted a hierarchical Bayesian, random effects analysis^41, 72^ for each subject. In this, the (suitably transformed) parameters **h**_*i*_ of experimental session *i* are treated as a random sample from a Gaussian distribution with means and variance ***θ*** = (***µ***_*θ*_, **Σ**_*θ*_}.

The prior distribution ***θ*** can be set as the maximum likelihood estimate:

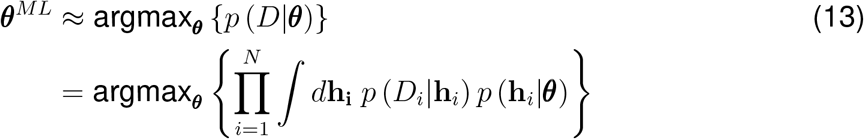

We optimized ***θ*** using an approximate Expectation-Maximization procedure. For the E-step of the k-th iteration, a Laplace approximation gives us

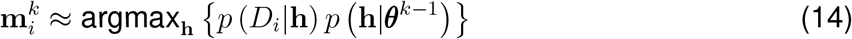

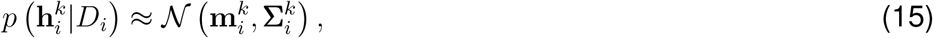

where 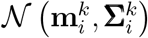 is the Normal distribution with the mean 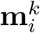 and the covariance 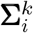 that is obtained from the inverse Hessian around 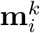. For the M step:

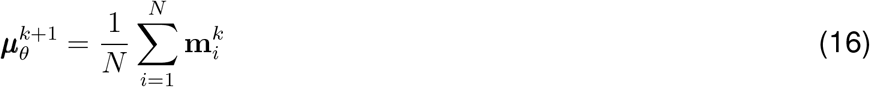

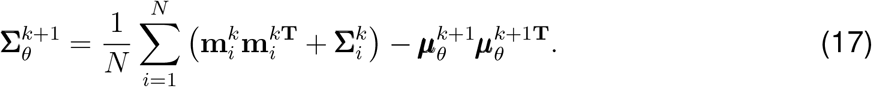

The maximum a posteriori probability (MAP) estimate for each subject *i* is

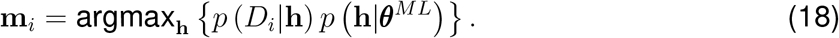

For simplicity, we assumed that the covariance 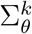 had zero off-diagonal terms, assuming that the effects were independent.

### Model comparison

We compared models according to their integrated Bayes Information Criterion (iBIC) scores.^41, 72^ We analyzed model log-likelihood log *p*(*D*|*M*):

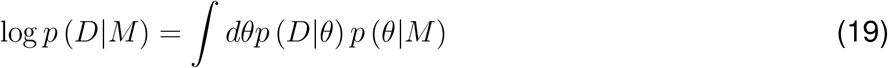

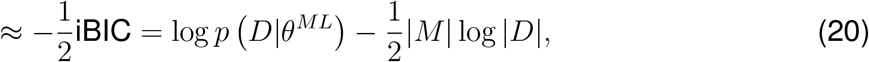

where iBIC is the *integrated* Bayesian Information Criterion, |*M* | is the number of fitted prior parameters and |*D*| is the number of data points (total number of choice made by all subjects). Here, log *p*(*D*|*θ*^*ML*^) can be computed by integrating out individual parameters:

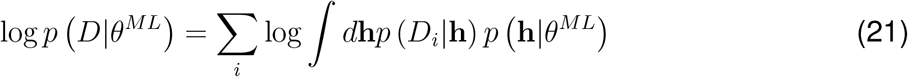

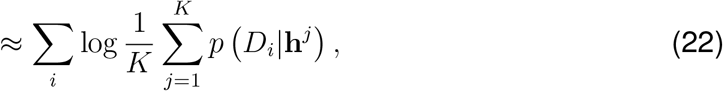

where we approximated the integral as the average over *K* samples **h**^*j*^’s generated from the prior *p* (**h**|*θ*^*ML*^).

### fMRI analysis

The data was processed using SPM 12. We performed a standard GLM analysis on the fMRI data with SPM 12. The SPM feature for asymmetrically orthogonalizing parametric regressors was disabled throughout. We first performed the analysis on all participants, performing the second-level inference pooled across all participants, and then HV and GD groups separately. The onsets of trial, action, and outcomes were controlled by stick regressors, separately for three trial types (gain, loss, neutral). In addition, the GLM had the following parametric regressors: at the trial onset, parametric modulators included the total fast value, the total slow value, and the output of the choice kernel, all separately for three trial types. At outcome onset, parametric regressors included the prediction error computed from fast value, the prediction error computed from slow value, and the outcome, all separated for three trial types. Movement regressors obtained from SPM preprocessing were also included.

### Validation analysis for between-group comparisons

To ensure that choosing ROIs on the basis of clusters of voxels identified as significant in the pooled group analysis for subsequent interrogation of group differences does not introduce a bias to produce erroneously significant results in the group difference analyses, we conducted a supplementary analysis, focusing on the slow value contrast as an example (but the simulation results should hold for all contrasts reported). We first calculated the covariance matrix between participants based on their beta values for the slow value over all grey matter voxels. We then generated synthetic beta values (10 million voxels for each participant) across participants, modeled as a multivariate Gaussian distribution with the real data’s mean and the calculated covariance. This synthetic data preserved the original correlation patterns observed between individuals and across groups.

Next, we applied a standard group-level analysis to identify which of the synthetic voxels exhibited significant correlations (large betas) by applying t-tests. Using *p <* 0.001 as a threshold, within these significant voxels, we estimated the likelihood of observing significantly different mean beta values between the HV and GD groups, with the significance threshold set at *p <* 0.05 by the permutation test, just like in our primary analysis.

We found that, across the synthetic voxels exhibiting ”significant” pooled group effects (at *p <* 0.001), the probability of subsequently finding a significant difference between the HV and GD groups in those voxels (tested by *p <* 0.05) was 0.036, confirming that the ROI selection did not favor a significant group difference over that which would be expected by chance. If we select the synthetic voxels that do not exhibit significant group-level effects (*p >* 0.2), the probability of finding a significant group difference was similar at 0.060. This analysis demonstrates that the ROI selection is not biasing the statistical inference conducted on those ROIs in favor of a significant group difference effect.

## Acknowledgement

We thank Ardy Rahman for implementing the recruitment and screening of participants, Jeff Cockburn, Caroline Charpentier, Vince Man, Reza Tadayon-Nejad, and Sandy Tanwisuth for valuable discussions. This work was supported by a grant from the International Center for Responsible Gaming to JOD and TF. KI is supported by the BBRF Young Investigator Grant.

